# Submerged plants may have substantial growth in shallow water lakes under nitrogen deposition: a theoretical investigation

**DOI:** 10.1101/019836

**Authors:** Ashehad A. Ali

## Abstract

1. Understanding shallow water lakes is important because they are not only subjected to a number of natural and anthropogenic stressors but are also able to switch from clear states with high levels of biodiversity to dominance by suspended sediments, phytoplankton, or floating plants. Such switches are often influenced by high levels of nitrogen deposition, which may reduce the diversity of submerged plant communities, suppressing benthic animal and plant life. Therefore, it is important to predict potential effects of nitrogen on such a change.
2. I modeled competition for light and nutrients between floating and submerged plants under low and high nutrient levels. Both types of plants were randomly chosen in their trait space by varying their average trait values by +/-25%.
3. Surprisingly, I found that submerged plants could perform better under high nitrogen by having substantial growth rates. Further, I found that responsiveness to nitrogen deposition is greater in shallow lakes with low biomass than high biomass. My results suggest that shallow lakes with low productivity may respond strongly to nitrogen deposition. Moreover, albedo may increase in the future over shallow lakes. Thus, future warming over shallow lakes may be higher than current predictions. Earth System Models should allow the trait values of submerged and floating plants to vary when making predictions about future climatic conditions.

## Introduction

Most of the world’s lakes are small and shallow (Wetzel, 1990), and many of them are subjected to a number of natural (Poff et al., 1997; Wilcox and Meeker, 1992) and anthropogenic stressors (Coops et al., 2003; Coops and Havens, 2005), such as nutrient deposition (Moss et al., 2013; Scheffer et al., 2003; Williamson et al., 2008). On the one hand, migratory fish, such as salmon, can bring nutrients up from marine environments into shallow water lakes (Naiman et al., 2002; Schindler et al., 2005). On the other hand, human-induced activities can cause increases in nitrogen deposition over shallow water lakes – as much as about a 30-fold increase in mean concentrations of nitrogen in lakes of Europe and North America has been documented (Bergstrom and Jannson, 2006).

Increased nitrogen deposition can result in eutrophication in shallow water lakes (Bergstrom and Jannson, 2006; Wolfe et al., 2001; Wolfe et al., 2002); where shifts in phytoplankton composition result in changes towards more mesotrophic species of diatoms (Bergstrom and Jannson, 2006; Scheffer et al., 1993) and increases in the organic matter deposited as sediment (Bergstrom and Jannson, 2006). Such shifts may reduce the diversity of submerged plant communities (Barker et al., 2008; James et al., 2005), leaving fewer niches for other animal and plant life (Jansen and Van Puijenbroek, 1998). Therefore, it is important to predict potential nitrogen effects on such change.

Submerged plants obtain much of their nutrient supply from sediments (Barko et al., 1991), but their growth is inhibited by low availability of light (Falkowski and Raven, 2007). Floating plant communities appear to be favoured by increased nutrient supply (Scheffer et al., 2003). Scheffer et al. (2003) suggested the establishment of floating plant dominance as a third, alternative stable state associated with high level of nitrogen deposition, the other two alternative stable states being submerged plant dominance and phytoplankton dominance.

However, experimental studies show that nutrient levels alone do not promote such a shift and suggest that an external driver is necessary (Balls et al., 1989; Irvine et al., 1989). Mesocosm experiments also reveal that nutrients alone are insufficient to displace a plant-dominated state (Feuchtmayr et al., 2007; Feuchtmayr et al., 2009). Warming can also affect the diversity of plants in shallow lakes (Mooij et al., 2005) but in this study I only consider nitrogen effects. Experimental data to date do not suggest a loss of submerged plant dominance through temperature increase alone, even at high rates of nitrogen deposition except in small temperate hypertrophic water bodies (Meerhoff et al., 2007).

In this study, I modeled competition for light and nutrients between floating and submerged plants and investigated shifts in plant community composition under low and high levels of nutrient deposition. To this end, I applied a probabilistic approach; in which, for both types of plants, parameter values for traits were sampled uniform randomly over reasonable ranges, and the model evaluated for each case. First, I identified the pattern of dominance of submerged and floating plants under high and low levels of nutrient deposition. Next, I identified suits of traits which are favoured by nitrogen deposition. Finally, I determined the biomass responsiveness of the submerged and floating plants to nitrogen deposition.

## Methodology

### Overview

I utilized the equations of Scheffer et al. (2003) to study the competition outcomes between submerged and floating plants under low and high nitrogen levels. Since the plant species of both types differ widely, different sets of parameter values would be needed to represent the possible different plant species each type. To account for this variation in species, we assumed average trait values for each vegetation type and randomly varied trait values around the average value by +/-25% and investigated the competition outcomes between submerged and floating plants for each set of trait values.

### Model

To model competition between submerged and floating plants under low and high nitrogen levels, I used the equations proposed by Scheffer et al. (2003) as given below;

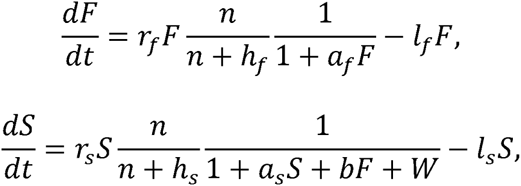

where changes over time of the biomass of floating plants, *F*, and submerged plants, *S,* are modeled as a function of their maximum growth rates, *r*_*f*_ and *r*_*s*_. Turnover rates ascribed to respiration and mortality for floating and submerged are denoted as *l*_*f*_, and *l*_*s*_, respectively. Nutrient limitation is a saturating function of the total inorganic nitrogen concentration, *n*, in the water column, which is assumed to be a decreasing function of plant biomass:

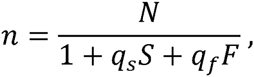

where the maximum concentration (*N*) in the absence of plants depends on the nutrient-loading of the system, and the parameters *q*_*s*_ and *q*_*f*_ represent the effect of submerged and floating plants on the nitrogen concentration in the water column. Light limitation is formulated as in Scheffer et al. (1997), where 1/*a*_*f*_ and 1/*a*_*s*_ are the densities of floating and submerged plants at which their growth rates become reduced by 50% because of intraspecific competition for light. In addition to this intraspecific competition, irradiation of submerged plants is reduced by light attenuation in the water column (*W*) and by shading by floating plants scaled by the parameter *b.* The parameter *h*_*s*_ is the half-saturation concentration at which nutrient supply from the sediment is sufficient to make submerged plant growth essentially independent of the nutrient concentration in the water column. The half-saturation concentration for floating plants is denoted as h_f_.

The default values and dimensions for parameters (traits) are given in Table 1. The default parameter values are chosen in such a way that they seem to mimic certain field situations in a reasonable way. Detailed description of the above model and its validation have been provided by Scheffer et al. (2003). Because plant species differ widely, different sets of parameter values are obviously needed to represent different plant species. I chose 10000 submerged and 10000 floating plants randomly in the trait space. Stochastic simulations for similar types of competition outcomes between alternative stable states has been performed elsewhere (e.g., 10000 simulations by Passarge et al. (2006)). I analyze how the results are affected by nitrogen deposition where plants compete for nutrient and light.

**Table 1.** Species traits used in the model, together with units and values used in model simulations. Trait values were taken from Scheffer et al. (2003). Mean trait values across the species were used as baseline values in the simulations. For the sensitivity analysis, the range of trait values was obtained by varying each trait by +/- 25%.

| **Trait** | **Definition** | **Baseline Trait Value**<br>**[Range]** | **Units** |
| --- | --- | --- | --- |
| $a_f$ | Parameter related to densities of floating plants at which their growth rates become reduced by 50% due to intraspecific light competition | 0.01 [0.0075,0.0125] | $(g\ dw\ m^{-2})^{-1}$ |
| $a_s$ | Parameter related to densities of submerged plants at which their growth rates become reduced by 50% due to intraspecific light competition | 0.01 [0.0075,0.0125] | $(g\ dw\ m^{-2})^{-1}$ |
| $b$ | Parameter related to shading by floating plants | 0.02 [0.0150,0.025,] | $(g\ dw\ m^{-2})^{-1}$ |
| $h_f$ | Half-saturation concentration for floating plants at which nutrient supply from the sediment is sufficient to make floating plant growth independent of the nutrient concentration | 0.2 [0.15,0.25] | $mg\ N\ liter^{-1}$ |
| $h_s$ | Half-saturation concentration for submerged plants at which nutrient supply from the sediment is sufficient to make submerged plant growth independent of the nutrient concentration | 0.0 [held constant] | $mg\ N\ liter^{-1}$ |
| $l_f$ | Turnover rates of floating plants due to respiration and mortality | 0.05 [0.0375,0.0625] | day<sup>-1</sup> |
| $l_s$ | Turnover rates of submerged plants due to respiration and mortality | 0.05 [0.0375,0.0625] | day<sup>-1</sup> |
| $q_s$ | Effect of submerged plants on the nitrogen concentration in the water column | 0.005 [0.0375,0.0625] | (g dw m<sup>-2</sup>)<sup>-1</sup> |
| $q_f$ | Effect of floating plants on the nitrogen concentration in the water column | 0.075<br>[0.05625,0.09375] | (g dw m<sup>-2</sup>)<sup>-1</sup> |
| $r_f$ | Maximum growth rates of floating plants | 0.5 [0.375,0.625] | day<sup>-1</sup> |
| $W$ | Water column | 0 [held constant] | unitless |
| $r_s$ | Maximum growth rates of submerged plants | 0.5 [0.375,0.625] | day<sup>-1</sup> |

## Results

### Substantial submerged growth under nitrogen fertilization

I found that nitrogen deposition reduced the dominance ratio of the winning plants by approximately 80% (Fig. 1), which is true for no matter which plant was dominant under low levels of nitrogen deposition. Submerged plants were found to win 75 percent of the time under low nitrogen (Fig. 1). In contrast, submerged plants won 25 percent of the time under nitrogen fertilization (Fig. 1). Based on the low nitrogen treatment, the model prediction is that the species that lose in competition has the proportion of biomass as 0.13% (Fig. 1), or equivocally winning plants dominated 98 percent of the time under low nitrogen (Fig. 1). I tested whether the number of times the plants lose under nitrogen deposition is more than 10% (as a threshold) (Table 2). The test reveals that the probability of dominance under nitrogen deposition is less than 0.9 (Table 2).

**Figure 1.**
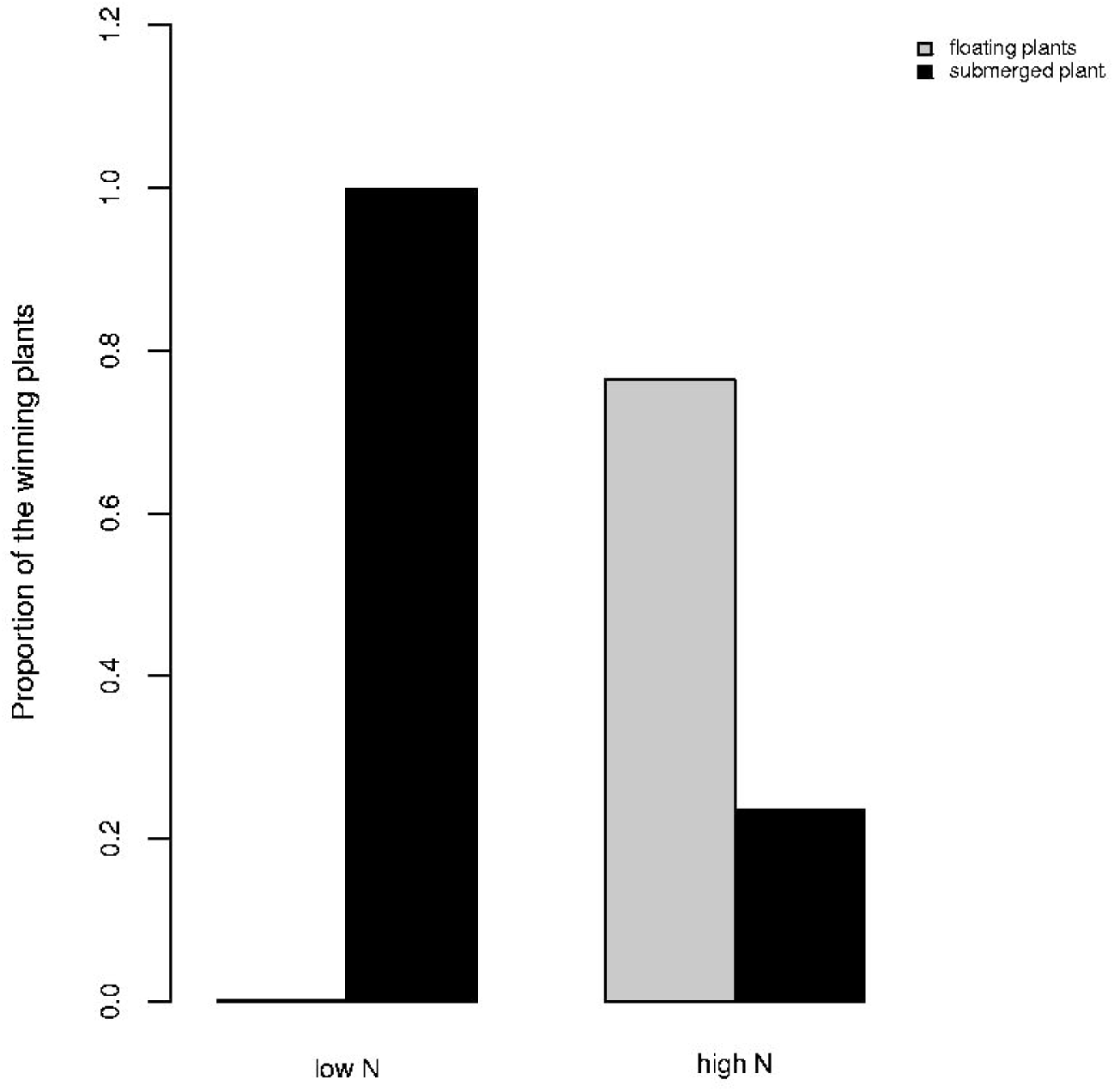
Proportion of winning plants (submerged or floating) at equilibrium under low and high N deposition. The proportion of winning plants was calculated as winners’ equilibrium biomass divided by the total biomass of the system at equilibrium.

**Table 2.** Testing the significance that the proportion of biomass of the losing species under high nitrogen will be more than 10%.

| Hypothesis | Z-score | p-value |
| --- | --- | --- |
| $H_0: \pi < 0.1$<br>$H_1: \pi \geq 0.1$ | 14.2935 | < 0.05 |

Figure 2 shows the biomass of the submerged plants under low nitrogen on the x-axis and its share of biomass under high nitrogen on the y-axis. All the points below the horizontal line at y = 0.5 indicate where the dominance of the submerged plants under low N changes at high N. There is a strong tendency for submerged plants with higher biomass under low N to have either low or high share of biomass under high N (Fig. 2). The conclusion is that N fertilization can favor submerged plants.

**Figure 2.**
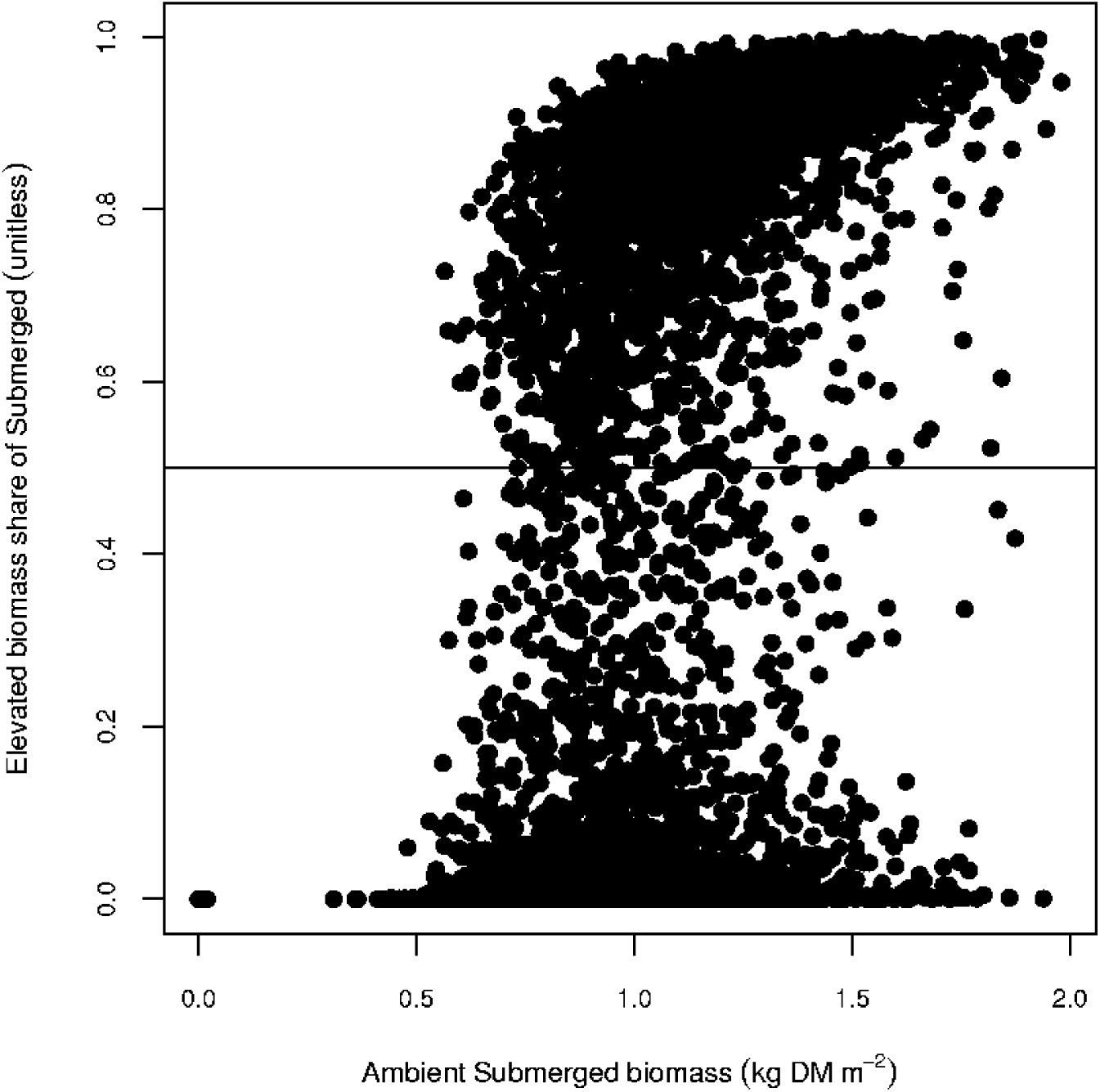
Biomass share of the submerged plants at high N as a function of its biomas (kg DM m^-2^) at low N. The horizontal line indicates where the biomass share of submerged plants at high N is 0.5.

### Suites of traits of submerged plants are favored by nitrogen deposition

Table 3 shows the mean trait values for the winning submerged and floating plants under low and high nitrogen deposition. The importance of each trait in determining the outcome of competition was evaluated by calculating the difference between the mean trait value of the winning submerged plants and winning floating plants, divided by the mean trait value overall. Under low nitrogen, the traits favouring success in competition were, in decreasing order of importance: parameter related to the effect of submerged plants on the nitrogen concentration in the water column (q_s_), parameter related to the effect of submerged plants on the nitrogen concentration in the water column (q_f_), parameter related to shading by floating plants (B), half-saturation concentration for floating plants at which nutrient supply from the sediment is sufficient to make floating plant growth independent of the nutrient concentration (h_f_), turnover rates of submerged plants due to respiration and mortality (l_s_), turnover rates of floating plants due to respiration and mortality (l_f_), parameter related to densities of submerged plants at which their growth rates become reduced by 50% due to intraspecific light competition (a_s_), parameter related to densities of floating plants at which their growth rates become reduced by 50% due to intraspecific light competition (a_f_), maximum growth rates of submerged plants (r_s_) and maximum growth rates of floating plants (r_f_) (Table 3). Nitrogen deposition highlighted a similar set of traits in determining the outcome of competition; however, r_s_ and r_f_ were more important than a_s_,a_f_,l_s_ and l_f_.

**Table 3.**
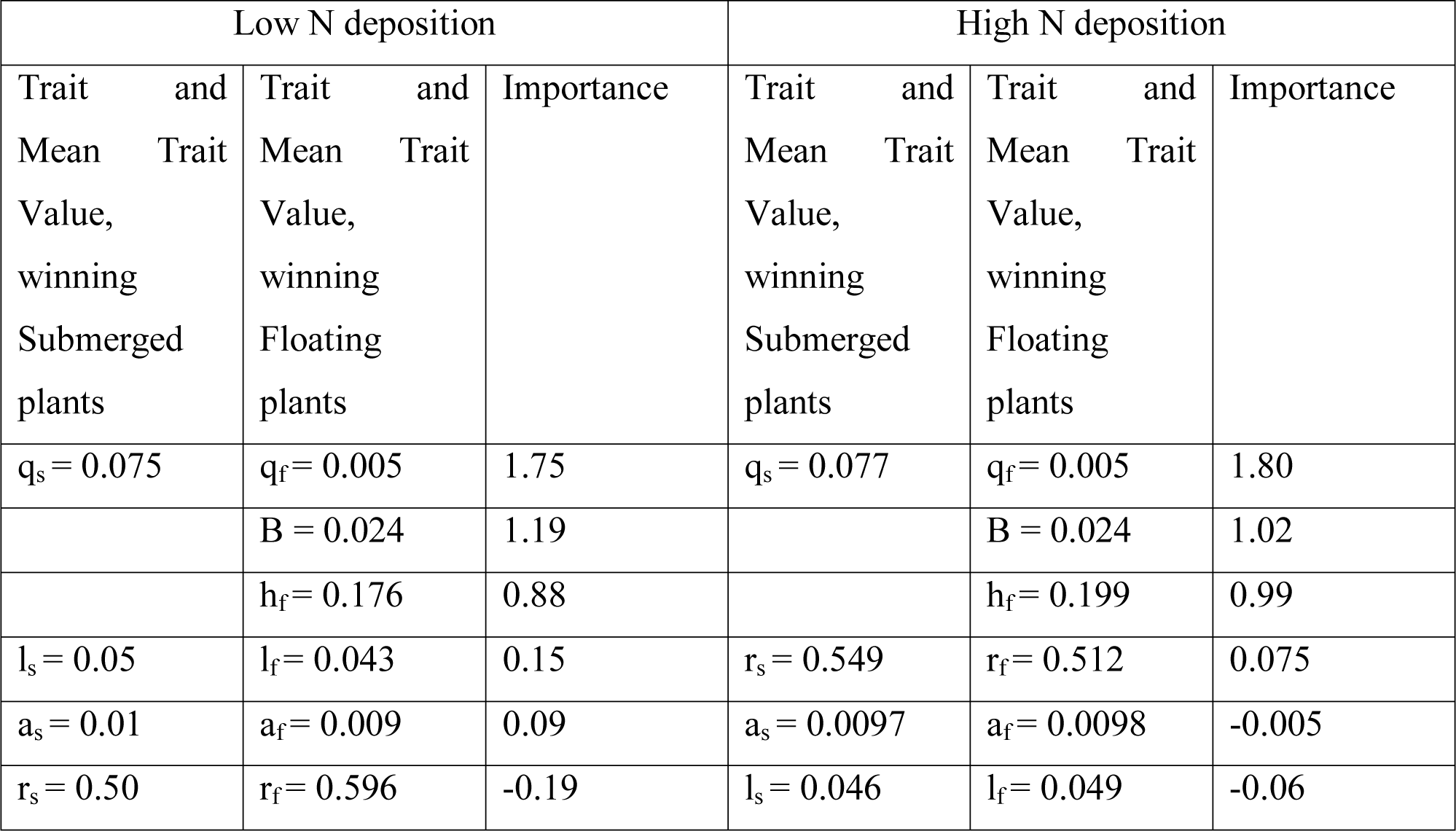
Comparison of mean trait values of winning submerged and winning floating plants when randomly-generated plants are compared under low and high N deposition. Importance of each trait in determining the outcome of competition is calculated as the difference between the average trait values of the winning submerged and winning floating plants, divided by the average trait value overall. Traits are ordered (in the descending order) by importance.

| Low N deposition |  |  | High N deposition |  |  |
| --- | --- | --- | --- | --- | --- |
| Trait and Mean Trait Value, winning Submerged plants | Trait and Mean Trait Value, winning Floating plants | Importance | Trait and Mean Trait Value, winning Submerged plants | Trait and Mean Trait Value, winning Floating plants | Importance |
| $q_s = 0.075$ | $q_f = 0.005$ | 1.75 | $q_s = 0.077$ | $q_f = 0.005$ | 1.80 |
| | $B = 0.024$ | 1.19 | | $B = 0.024$ | 1.02 |
| | $h_f = 0.176$ | 0.88 | | $h_f = 0.199$ | 0.99 |
| $l_s = 0.05$ | $l_f = 0.043$ | 0.15 | $r_s = 0.549$ | $r_f = 0.512$ | 0.075 |
| $a_s = 0.01$ | $a_f = 0.009$ | 0.09 | $a_s = 0.0097$ | $a_f = 0.0098$ | -0.005 |
| $r_s = 0.50$ | $r_f = 0.596$ | -0.19 | $l_s = 0.046$ | $l_f = 0.049$ | -0.06 |

### Biomass responsiveness to nitrogen deposition

Since submerged plants were found to win 75 percent of the time under low nitrogen (Fig. 1), by definition, submerged plants had higher biomass than floating plants. I found that responsiveness to nitrogen deposition was greater in shallow lakes with low biomass than high biomass (Fig. 3). Submerged plants responded less strongly to nitrogen deposition than floating plants (Fig. 3).

**Figure 3.**
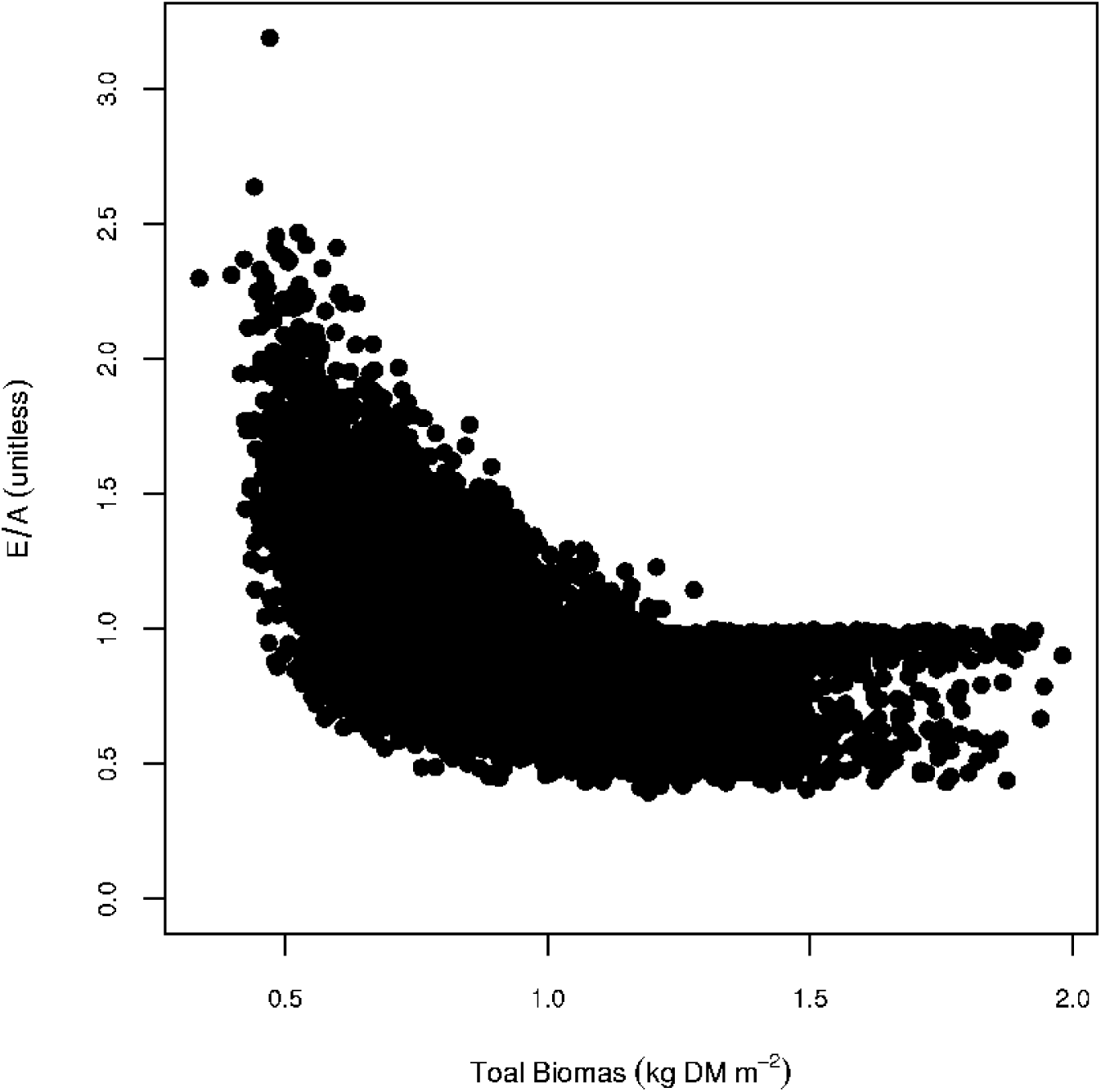
Enhancement ratio (E/A) of total biomass of the system (equilibrium biomass at high N divided by that at ambient N) as a function of equilibrium total biomass at ambient N, where the total biomass is the sum of biomasses of submerged and floating plants. The highest biomass enhancement ratios are obtained for the lowest ambient biomass.

## Discussion

### Summary

The model predicts that nitrogen deposition would reduce the dominance ratio of the winning species. Further, submerged plants may have substantial growth under nitrogen fertilization. The model also predicts that responsiveness to nitrogen deposition would be greater in shallow lakes with low biomass than those with high biomass. This study provides an innovative, mechanistic hypothesis for the outcomes of competition under nitrogen fertilization that can be tested experimentally. This is also the first systematic modelling study exploring how submerged and floating plants are likely to vary in response to nitrogen deposition based on the suite of traits they possess.

### Substantial submerged growth under nitrogen fertilization

Certain traits of the submerged plants may help them to benefit from increased nutrient availability (Scheffer et al., 2003). In this work, nitrogen fertilization tended to favor submerged plants with maximum growth rates. Low light and high nutrient availability has been shown to alter growth and allocation patterns in a small scale experiment on macrophytes (Cronin and Lodge, 2003). This study showed that submerged plants will not necessarily be replaced by floating plants at nitrogen deposition. This theoretical finding is in line with an experiment on mesocosms (Feuchtmayr et al., 2009), where they varied nitrogen. In this study, variation of the traits around the average value by +/-25% yielded 25% dominance of submerged plants under N fertilization. In reality, traits of plants could vary by a larger amount (e.g. by +/-50% of the mean value) (Ali et al., 2013). Therefore, the dominance of submerged plants under N fertilization could be even larger. This suggests that albedo may increase in future over shallow lakes. Thus, future warming over shallow lakes may be higher than current predictions. Earth System Models (ESMs) should allow the trait values of submerged and floating plants to vary when making predictions about future climatic conditions. My suggestion for ESMs in this regard is in line with a study on terrestrial plants (Verheijen et al., 2012), who illustrated that allowing plant traits within plant functional types to vary with plant trait - climate relationships yielded a closer a match to some types of observational data. I posit that traits of submerged and floating plants could also evolve through time, which will have important consequences for modeling dynamics of shallow water lakes.

### Biomass responsiveness to nitrogen deposition

I have shown that there are considerable differences among hypothetical trait-simulated submerged and floating plants in responsiveness to nitrogen deposition. Submerged plants responded less strongly to nitrogen deposition than floating plants because at low levels of nitrogen deposition, submerged plants are only light limited whereas floating plants are nutrient limited (Gaudet and Muthuri, 1981; Weisner et al., 1997). When nitrogen is added, floating plants are relieved of nutrient limitation and absorb a greater proportion of light, which enhances their biomass. An increase in the biomass of floating plants increases light attenuation in the water column greatly and so submerged plants respond little to nitrogen deposition.

### Caveats

The results of my study apply to situations in which case dominance by submerged plants is not prevented by factors other than nutrients. Competition among submerged and floating plants was examined at equilibrium – the initial responses do not seem to matter too much, but if the outcome of competition is decided at establishment of the plants then equilibrium responses are not helpful. The traits were treated independent, when in fact they are likely to be correlated.

### Potential mechanisms that were not considered

I did not consider a number of mechanisms that might regulate the abundance of submerged and floating plants. One mechanism is related to fluctuations in water levels. High water levels have been reported to be quite important in nutrient regeneration from mud (Gaudet and Muthuri, 1981), whereby large reserves of nutrients get released to the overlying water. In contrast, lower water level may favor submerged plants (Blindow et al., 1993; Lauridsen et al., 1994). Another mechanism links fluctuations in submerged plants abundance to population changes in grazers (Bales et al., 1993). In one field enclosure experiment, positive effects on macrophyte growth after reduction of fish predation on ephiphyte-grazing snails were observed (Martin et al., 1992; Underwood et al., 1992). A third mechanism is about the release of allelopathetically active substances, which may cause the negative effect of submerged plants on floating (Blindow, 1992; Scheffer et al., 1993); however, proof for allelopathy *in situ* is difficult or even inaccessible (Gross et al., 2007).

## Acknowledgements

This work is funded by UC Lab Research Program (Award ID: 2012UCLRP0IT00000068990) and the DOE Office of Science Next Generation Ecosystem Experiment at Artic (NGEE-Arctic) Program.

